# DHCR24-mediated sterol homeostasis during spermatogenesis is required for sperm mitochondrial sheath formation and impacts male fertility over time

**DOI:** 10.1101/2023.12.21.572851

**Authors:** Sona Relovska, Huafeng Wang, Xinbo Zhang, Pablo Fernández-Tussy, Kyung Jo Jeong, Jungmin Choi, Yajaira Suárez, Jeffrey G. McDonald, Carlos Fernández-Hernando, Jean-Ju Chung

## Abstract

Desmosterol and cholesterol are essential lipid components of the sperm plasma membrane. Cholesterol efflux is required for capacitation, a process through which sperm acquire fertilizing ability. In this study, using a transgenic mouse model overexpressing 24-dehydrocholesterol reductase (DHCR24), an enzyme in the sterol biosynthesis pathway responsible for the conversion of desmosterol to cholesterol, we show that disruption of sterol homeostasis during spermatogenesis led to defective sperm morphology characterized by incomplete mitochondrial packing in the midpiece, reduced sperm count and motility, and a decline in male fertility with increasing paternal age, without changes in body fat composition. Sperm depleted of desmosterol exhibit inefficiency in the acrosome reaction, metabolic dysfunction, and an inability to fertilize the egg. These findings provide molecular insights into sterol homeostasis for sperm capacitation and its impact on male fertility.

## Introduction

Lipids are essential for mammalian fertility, contributing to both steroidogenesis (Miller and Auchus, 2011) and germ cell formation (Shi et al., 2018). In humans and other mammalian species, cholesterol and desmosterol are major sterol components of the sperm plasma membrane (Keber et al., 2013). To gain fertilization capability, the sperm plasma membrane undergoes changes that result in cholesterol efflux, allowing sperm to undergo capacitation, including the acrosome reaction (Leahy and Gadella, 2015). An appropriate species-dependent sterol composition of the sperm plasma membrane is essential for successful fertilization. Coincidentally, male obesity has been linked to reduced reproductive success and infertility in both humans (Leisegang et al., 2021, Craig et al., 2017) and animal models (Ghanayem et al., 2010; Borges et al., 2017; Han et al., 2023). During spermiation, when mature spermatids dissociate from Sertoli cells and enter the lumen of the seminiferous tubules, the cholesterol content of the sperm plasma membrane decreases. This reduction continues as sperm migrate through the epididymis and acquire fertilizing capacity (Nikolopoulou et al., 1985; Haidl and Opper, 1997). During sperm capacitation, the plasma membrane becomes more fluid, and cholesterol relocates to the apical margin of the sperm head before eventually translocated to the extracellular environment by albumin (Leahy and Gadella, 2015). A study in boar and mouse sperm demonstrated the importance of sterol oxidation and oxysterol formation, which is dependent on the bicarbonate-induced reactive oxygen species (ROS) signaling pathway, for sterol removal from the plasma membrane during capacitation (Boerke et al., 2013). In sperm samples from infertile men, elevated levels of both cholesterol and its immediate precursor desmosterol were observed, but the cholesterol/desmosterol ratio was decreased (Zalata et al., 2010), suggesting that sperm sterol homeostasis is essential for male fertility in humans. However, the specific impact of the disruption of sterol homeostasis on sperm physiology remains largely unknown.

Two parallel pathways contribute to cellular cholesterol biosynthesis from lanosterol. Both pathways – the Bloch pathway and the Kandutsch-Russell pathways – are interconnected by the enzyme DHCR24, which reduces the double bond in the side chain of all Bloch pathway sterols and converts them to the Kandutsch-Russel pathway sterols (Figure 1A). Notably, DHCR24 activity in various tissues is essential for the production of diverse bioactive sterols, crucial for maintaining cellular homeostasis and tissue physiology. The *Dhcr24*-null mouse model exhibits nearly undetectable levels of plasma cholesterol and abnormally high levels of desmosterol, leading to increased prenatal mortality, poor growth in males that persists into adulthood, degenerated testes, and infertility in both males and females (Wechsler et al., 2003). In humans, mutations in the *DHCR24* gene cause desmosterolosis, a rare congenital disorder characterized by excessive desmosterol and insufficient cholesterol. This condition also affects brain and head development and causes cardiac malformations and arthrogryposis (FitzPatrick et al., 1998; Andersson et al., 2002; Schaaf et al., 2011; Zolotushko et al., 2011; Dias et al., 2014; Rohanizadegan & Sacharow, 2018; Hill et al., 2023). Our previous studies demonstrated that DHCR24 overexpression in myeloid cells depletes desmosterol, increases inflammation and mitochondrial ROS production, and impairs LXR/RXR pathway activation (Zhang et al., 2021^a^).

**Figure 1.**
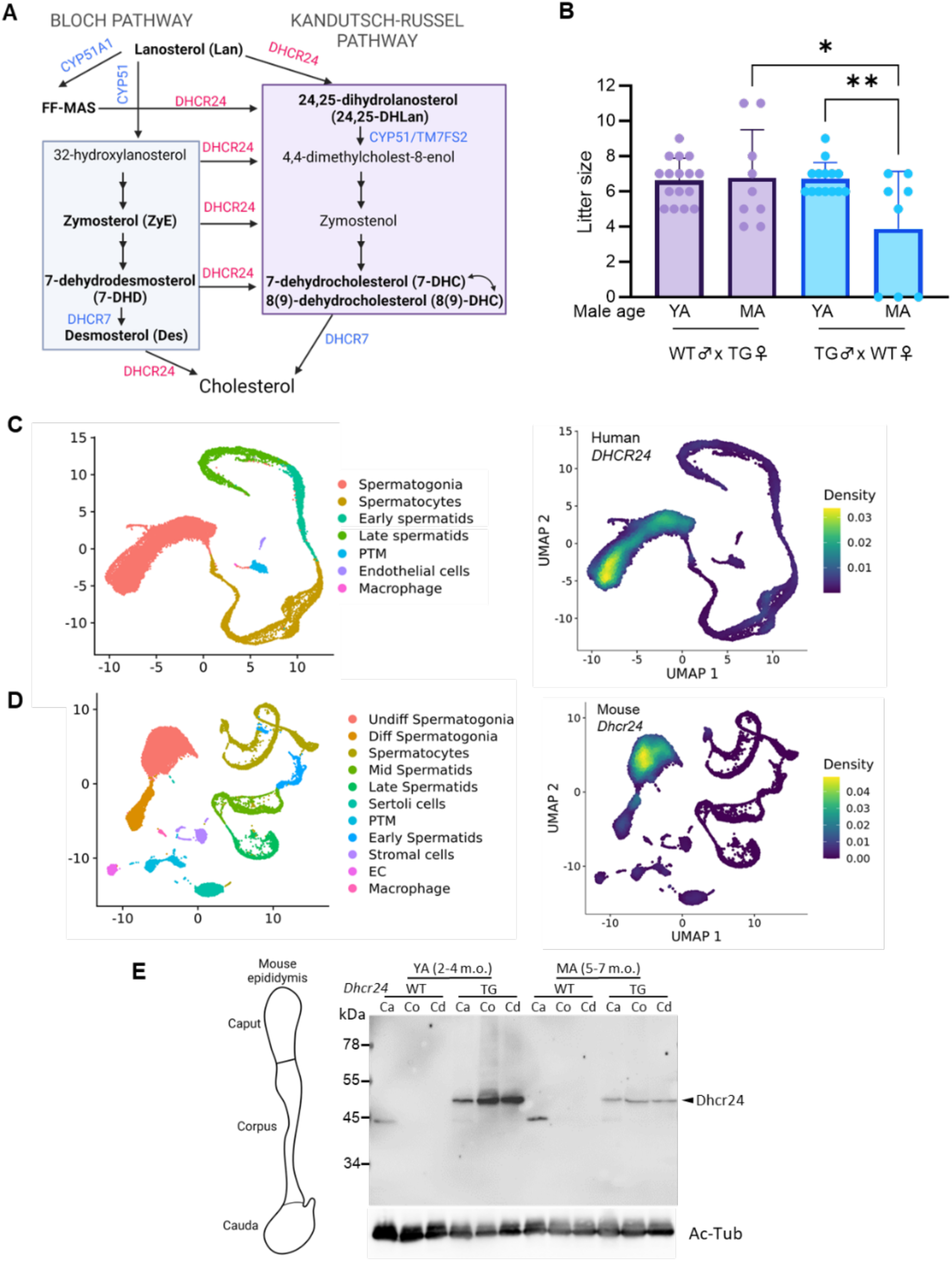
DHCR24 overexpression leads to age-dependent decline in male fertility. (**A**) Diagram showing the involvement of DHCR24 in two parallel pathways to produce cholesterol: the Bloch (*left*) and Kandutsch-Russel (*right*) pathways. (**B**) Litter sizes resulting from wild-type (WT) and *Dhcr24-TG* (TG) crosses of young adult (YA) and mature adult (MA) mice. Error bars SD, statistical analysis one-way ANOVA. (**C-D**) *Dhcr24* mRNA expression in human (C) and mouse (D) testicular cells using human (GSE109037) and mouse (GSE109033) testis single-cell RNA (scRNA)-seq datasets. UMAP plots represent 7 and 11 clusters of testicular cells in human and mouse (*left*) and nebula plots of *Dhcr24* mRNA expression density in these clusters (*right*). EC, endothelial cell; PTM, peritubular myoid cell. (**E**) Western blot analysis of DHCR24 protein levels in epididymal spermatozoa from caput (Ca), corpus (Co), or cauda (Cd) of WT and *Dhcr24-TG* young and mature adults. YA, young adult (2-4 months old); MA, mature adult (5-7 months old). Illustration created with BioRender.com.

In this study, we report that whole-body overexpression of DHCR24 (*Dhcr24-TG*) leads to decreased fertility with reduced sperm count and motility in male mice, correlated with increasing paternal age. DHCR24 overexpression depletes sperm desmosterol and disrupts plasma membrane sterol homeostasis. Scanning and transmission electron microscopy of *Dhcr24-TG* spermatozoa revealed defects in the mitochondria and sperm tail, including incomplete mitochondrial packing and bent tails. IZUMO staining showed reduced efficiency of *Dhcr24-TG* sperm to undergo the acrosome reaction during capacitation, even in spermatozoa from young adult males. Furthermore, our findings suggest that metabolically, uncapacitated *Dhcr24-TG* sperm resemble capacitated sperm in their increased oxygen consumption and faster depolarization of mitochondrial membrane potential. Taken together, our results demonstrate that desmosterol depletion and/or altered sterol homeostasis during spermatogenesis impacts sperm morphology, number, motility, and metabolism, leading to reduced male fertility.

## Results

### DHCR24 overexpression leads to an age-dependent decline in male fertility

In our previous study, we generated a myeloid-specific conditional knock-in mouse line overexpressing DHCR24 (Figure 1A) (Zhang et al., 2021^a^). Here, we used a similar approach to generate a global *Dhcr24* transgenic mouse line by breeding the *Dhcr24^fl/fl^*mice (overexpression of the construct in the *Rosa26* genetic locus) with the *EIla^CRE^*mice (which express *Cre recombinase* in the early mouse embryo) (Figure 1–figure Supplement 1A). We used male mice aged 2 to 10 months, categorizing them as young (2-4 months old), mature (5-7 months old) and middle-aged (8-10 months old) adults (adapted from Flurkey et al., 2007). After establishing mating between wild-type (WT) and *Dhcr24-TG* (TG) animals, using young and mature adult males and young adult females, we found that mature adult TG males sired significantly smaller litters than young adult TG males, with no alterations in the pup sex ratio (Figure 1B, Figure 1– figure supplement 1B). Moreover, they exhibited delay in impregnating the females (Figure 1–figure Supplement 1C). In contrast, WT males displayed no fertility changes within this age range. In light of these observations, we delved into exploring the consequences of desmosterol depletion through DHCR24 overexpression on sperm physiology.

Analyzing publicly available human (GSE109037) and mouse (GSE109033) testis scRNA-seq data, we found that *DHCR24* mRNA in human testis is most abundantly expressed in spermatogonia and, to a lesser extent, in spermatocytes and late spermatids (Figure 1C). This pattern mirrors *Dhcr24* mRNA levels in mouse adult and postnatal day 6 (P6) testis (Figure 1D), where it is most abundant in spermatogonia and to a lesser extent in Sertoli cells and peritubular myoid cells. We examined DHCR24 protein expression in WT and TG testes and found increased protein level in samples from TG animals (Figure 1–figure Supplement 1D). In line with the scRNA-seq data, DHCR24 is not readily detected in WT spermatozoa, but is detected in young adult TG spermatozoa from caput, corpus and cauda epididymis, and to a lesser extent in mature adult TG spermatozoa (Figure 1E). Collectively, these data demonstrate DHCR24 expression in male germ cells and suggest that disruption of sterol synthesis during spermatogenesis may explain the observed decline of male fertility in *Dhcr24-TG* animals.

### DHCR24 overexpression leads to dysregulation of sperm sterol composition and capacitation efficiency

Alterations in sperm cholesterol, desmosterol, and the cholesterol to desmosterol ratio have been associated with subfertility in men (Zalata et al., 2010). To assess changes in sperm sterol composition, we measured specific sterols in the Bloch and Kandutsch-Russel pathways in spermatozoa from mature adult WT and TG males. In the Bloch pathway, we observed decreased levels of desmosterol (Des), 7-dehydrodesmosterol (7-DHD), and zymosterol (ZyE) (Figure 2A). In the Kandutsch-Russell pathway, both immediate cholesterol precursors –7-dehydrocholesterol (7-DHC) and 8,9-dehydrocholesterol (8-DHC)– showed decreased levels compared to WT sperm. Notably, the levels of 24,25-dihydrolanosterol (24,25DHLan) were significantly increased albeit just above the detection range (Figure 2B). Although we did not directly measure sperm cholesterol by the mass spectrometry, we speculated that cholesterol might be increased in TG sperm, considering the predominant flux (97%) through the Bloch pathway in the testis (Mitsche et al., 2015). However, our results indicated no statistical difference in cholesterol levels isolated from the whole testis of young adult and mature adult WT or Dhcr24-TG males (Figure 2C). This is consistent with our findings of no significant changes in mRNA expression in *Abca1* cholesterol transporter in the testis of young adult and mature adult WT and *Dhcr24-TG* males, nor *Nr1h2* and *Nr1h3,* which encode the ABCA1 transcriptional regulators, LXRβ and LXRα, respectively, (Figure 2–figure Supplement 2A). Additionally, the ABCA1 protein levels remained unchanged (Figure 2–figure Supplement 2B). Furthermore, EchoMRI results showed no changes in the fat mass to lean mass ratio in middle-aged TG males (Figure 2–figure Supplement 3A), consistent with unchanged plasma cholesterol levels between young adult WT and *Dhcr24-TG* males (Figure 2–figure Supplement 3B), further supporting these findings.

**Figure 2.**
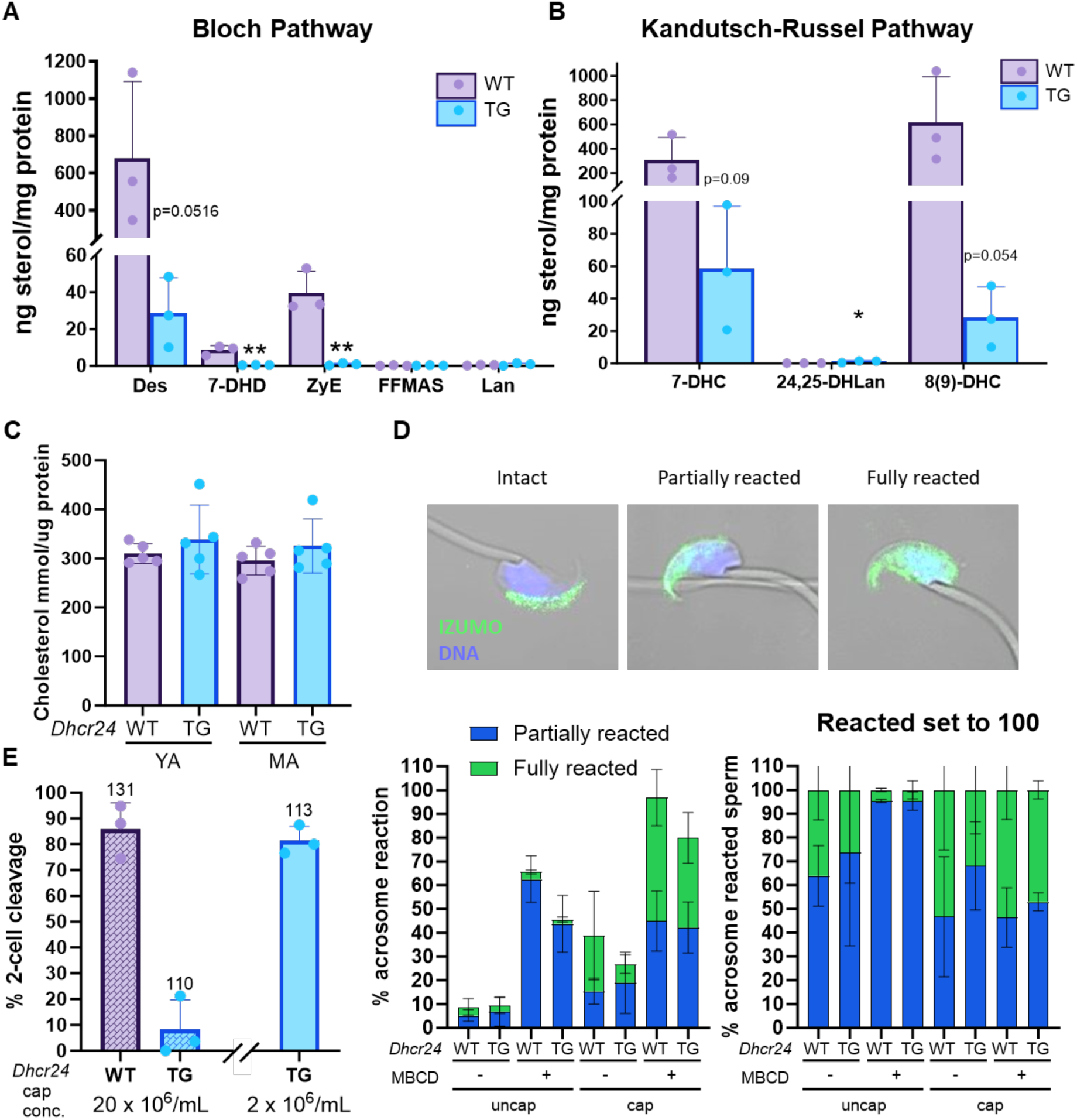
Dysregulated sterol biosynthesis affects sperm plasma membrane of *Dhcr24-TG* males. (**A** and **B**) Lipidomics analysis of WT and *Dhcr24-TG* (TG) sperm from mature adult mice for lipid species in Bloch (A) and Kandutsch-Russel (B) pathways. Statistical analysis student’s t-test. (**C**) Cholesterol levels from the testis of WT or *Dhcr24-TG* young adult and mature adult animals. (**D**) Acrosome reaction progression by IZUMO staining. Spermatozoa from young adult WT and *Dhcr24-TG* (TG) animals were incubated under non-capacitating (uncap) or capacitating (cap) conditions in the absence (-) or presence (+) of methyl-β-cyclodextrin (MBCD). (**E**) *In vitro* fertilization using spermatozoa of WT and *Dhcr24-TG* (TG) young adult animals capacitated at high (∼20×10^6^ sperm/mL) or standard (2×10^6^ sperm/mL) cell concentration. All error bars SD.

To determine whether the altered sterol composition of TG sperm affects their ability to undergo the acrosome reaction, we incubated them in the absence or presence of bovine serum albumin (BSA) or methyl-β-cyclodextrin (MBCD), both acting as a cholesterol sink during capacitation, with MBCD even more efficiently than BSA (Visconti et al., 1999). As cholesterol efflux is required for the acrosome reaction to occur in a timely manner (Leahy and Gadella, 2015), fixed spermatozoa were immunostained for IZUMO to examine the status of acrosome reaction. We found that TG spermatozoa did not undergo acrosome reaction as efficiently as WT spermatozoa, *i.e.,* the proportion of partially reacted TG spermatozoa was higher (Figure 2D). These observations align with the experiment using spermatozoa from middle-aged adult animals, where a greater proportion of TG spermatozoa were only partially acrosome-reacted than WT spermatozoa (Figure 2–figure Supplement 3C). Notably, even sperm from young adult TG males demonstrated reduced efficiency during *in vitro* fertilization (IVF). When TG sperm were capacitated at a high concentration (20×10^6^ cells/mL), the 2-cell cleavage rate dropped below 10%, whereas WT sperm still fertilized the oocytes with over 80% cleavage rate (Figure 2E). In contrast, TG sperm capacitated at the standard concentration used for IVF (2×10^6^ cells/mL) fertilized the oocytes at 80%. Taken together, these results demonstrate that altered sterol composition affects the efficiency of sperm capacitation and impairs fertility.

### *Dhcr24-TG* males have decreased sperm count and motility

As DHCR24 is globally overexpressed in our TG animals and not specifically in male germ cells, we investigated whether the changes in sterol composition have an impact on spermatogenesis and/or sperm motility. We found a lower epididymal sperm count in young adult TG males, although not statistically significant, but this difference became statistically significant with aging (Figure 3A). Consistently, hematoxylin and eosin staining revealed an overall normal testis histology, but mature adult TG testis and epididymis showed a decrease in sperm count (Figure 3B). To assess sperm total and hyperactivated motility, we utilized the Computer Assisted Sperm Analyzer (CASA) on WT and TG spermatozoa. Both before and after capacitation, spermatozoa from either young or mature adult TG animals displayed reduced motility. Additionally, the hyperactivated motility of TG spermatozoa was also diminished (Figure 3C). Similar results were observed in middle-aged adult spermatozoa capacitated in the presence of BSA or MBCD (Figure 3–figure Supplement 4A and 4B). Notably, the percentage of hyperactivated WT spermatozoa decreased when capacitated at high concentration (5.1%, compared to 16% percent when capacitated at the standard concentration), and the percentage of hyperactivated TG sperm was 3-5 times lower than that of WT, regardless of sperm concentration (1.7% and 2.9% at high and standard concentrations, respectively) (Figure 3–figure Supplement 4D). Importantly, sperm concentration during capacitation did not affect the total motility of either WT or TG spermatozoa (Figure 3–figure Supplement 4C). Flagellar waveform analysis revealed that a fraction of TG spermatozoa exhibited altered flagellar beating that became more prominent during sperm capacitation, presumably due to defective flagellar bending that flips the head backward (Figure 3D, Figure 3–supplementary video 1). These motility defects prompted us to further investigate sperm morphology in more detail.

**Figure 3.**
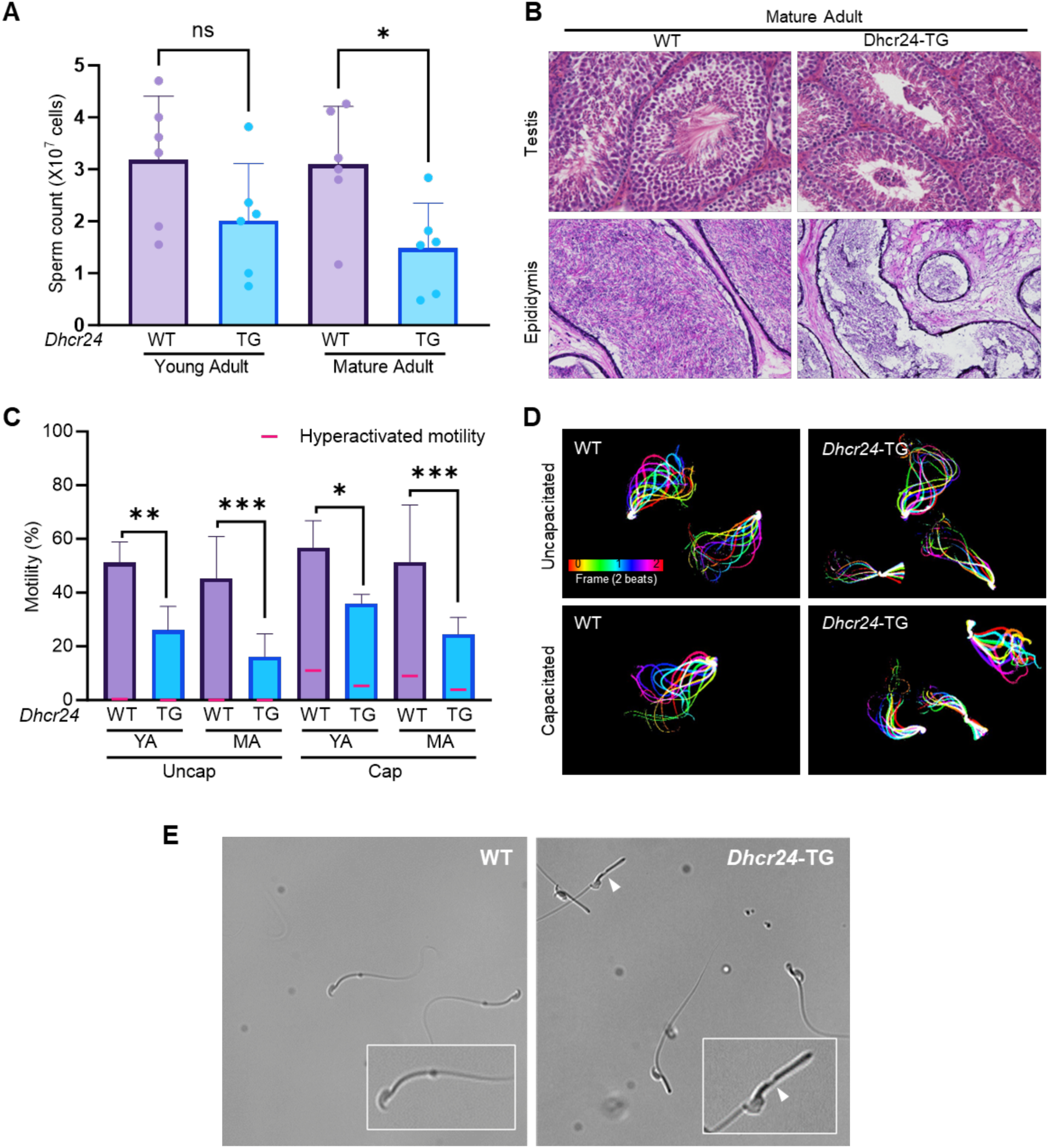
*Dhcr24-TG* males have decreased sperm count and motility. (**A**) Sperm counts from young and mature adult WT and *Dhcr24-TG* (TG) male mice. Statistical analysis one way ANOVA. (**B**) Hematoxylin and eosin staining of testis and epididymis from mature adult WT and *Dhcr24-TG* mice. (**C**) Total and hyperactivated motility from spermatozoa of young and mature adult WT and *Dhcr24-TG* males before and after capacitation (n=4). Statistical analysis two-way ANOVA. (**D**) Flagellar waveform analysis of WT and *Dhcr24-TG* spermatozoa before and after capacitation (young adult). (**E**) Brightfield images of spermatozoa from young adult WT and *Dhcr24-TG* animals. White arrow points to thinner area in the flagellum in *Dhcr24-TG* sperm. YA, young adult (2 – 4 months old); MA, mature adult (5 – 7 months old). All error bars SD.

### Desmosterol depletion results in sperm tail bending and aberrant mitochondrial packing

Upon closer examination of our movies, we found that bent TG spermatozoa often exhibited a head curled toward the midpiece and a thinning in the distal midpiece just before the annulus (Figure 3E, Figure 3– supplementary video 1). Co-immunostaining for the outer dense fiber (ODF) component ODF2 and the mitochondrial protein Tom20 indicated defective mitochondrial packing and ODF exposure (Figure 4A). Further examination of TG sperm ultrastructure using scanning electron microscopy (SEM) and transmission electron microscopy (TEM) (Figure 4B and 4D) revealed the inverted heads and thinning of the distal midpiece of TG spermatozoa, which lacked the mitochondrial sheath and exposed the outer dense fiber due to mitochondrial sheath shortening (Figure 4B and 4C). Quantification and classification of mitochondrial unraveling and tail bending defects from the SEM images confirmed the increased instances of aberrant mitochondrial packing and defective structural integrity of TG spermatozoa (Figure 4D). TEM images further supported these findings, revealing defective mitochondrial structure and packing along the sperm tails in TG spermatozoa (Figure 4D).

**Figure 4.**
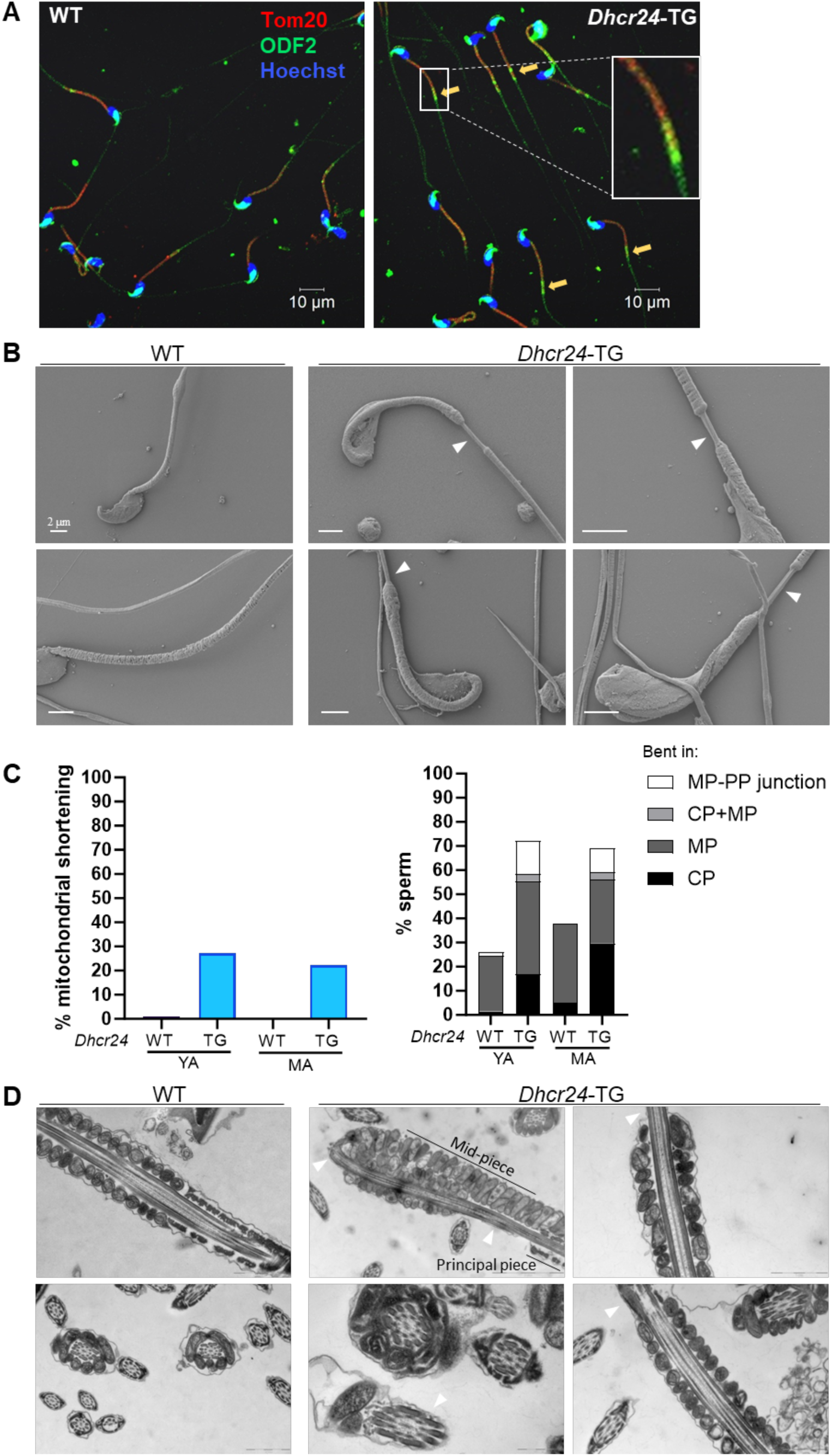
Sperm from *Dhcr24-TG* mice show abnormal head bending and incomplete mitochondrial packing. (**A**) Immunofluorescence images of WT and *Dhcr24-TG* spermatozoa from young adult animals stained with antibodies against Tom20 (red), ODF (green) and counterstained with Hoechst detecting nucleus (blue). (**B**) Scanning electron microscopy (SEM) images of WT and *Dhcr24-TG* spermatozoa from young (WT – bottom, *Dhcr24-TG* – top right, bottom two) and mature adult (WT – top, *Dhcr24-TG* – top left) animals. The bending of the head towards the midpiece can be observed in *Dhcr24-TG* spermatozoa. White arrows indicate the exposed outer dense fiber (ODF) in the midpiece of *Dhcr24-TG* spermatozoa. (**C**) Quantification of sperm defects in SEM images of sperm from young and mature adult WT and *Dhcr24-TG* animals. *Left*, percentage of sperm with shortened mitochondrial sheath; *right*, percentage of sperm with tail bending at the connecting piece (CP), midpiece (MP), both connecting piece and midpiece (CP+MP), or at the junction of midpiece-principal piece (MP-PP). (**D**) Transmission Electron Microscopy (TEM) images of WT and *Dhcr24-TG* spermatozoa from young (WT – top) and mature adult (WT – bottom, *Dhcr24-TG* – all) animals. White arrows indicate the exposed outer dense fiber (ODF) in the midpiece of *Dhcr24-TG* spermatozoa. The erratic mitochondrial packing along the sperm tail results in ODF exposure in the distal region of *Dhcr24-TG* spermatozoa. YA, young adult (2 – 4 months old); MA, mature adult (5 – 7 months old).

### Defective mitochondria in *Dhcr24-TG* males cause alteration in sperm metabolism

Having observed defects in mitochondrial structure in TG sperm, we next set out to investigate mitochondrial function. Spermatozoa must meet high energy demands as they travel to deliver the paternal genome to the oocyte. ATP production in spermatozoa occurs through two spatially compartmentalized metabolic processes in the tail: mitochondrial oxidative phosphorylation (OXPHOS) in the midpiece and glycolysis from the principal piece (Visconti, 2012). We measured sperm mitochondrial membrane potential (ΔΨm) using MitoTracker Red CMXRos dye (Pendergrass et al., 2004; Santiani et al., 2016), followed by antimycin treatment (Figure 5A), as previously described (Ferreira et al., 2021; Wang et al., 2022). We found that uncapacitated TG sperm from young adult males exhibited similar ΔΨm depolarization to capacitated WT and TG sperm after antimycin treatment – faster and to a greater extent than that of uncapacitated WT sperm (Figure 5B). This suggests a metabolic similarity of uncapacitated TG sperm to capacitated sperm. There was no difference in the rate or amplitude of membrane depolarization of the capacitated spermatozoa. To investigate whether this change affects oxygen consumption rate (OCR), we measured extracellular flux using Seahorse. Basal respiratory OCR, the difference between the initial and post-antimycin/rotenone injection measurements, increased after incubation under capacitating conditions, consistent with previous reports (Balbach et al., 2020, Ferreira et al., 2021). Interestingly, TG sperm from middle-aged males exhibited higher basal respiratory OCR in both uncapacitated and capacitated conditions (Figure 5C; Figure 5–figure Supplement 5), suggesting that uncapacitated TG spermatozoa have higher metabolic demands and are more comparable to capacitated spermatozoa, in line with our observations from ΔΨm measurements.

**Figure 5.**
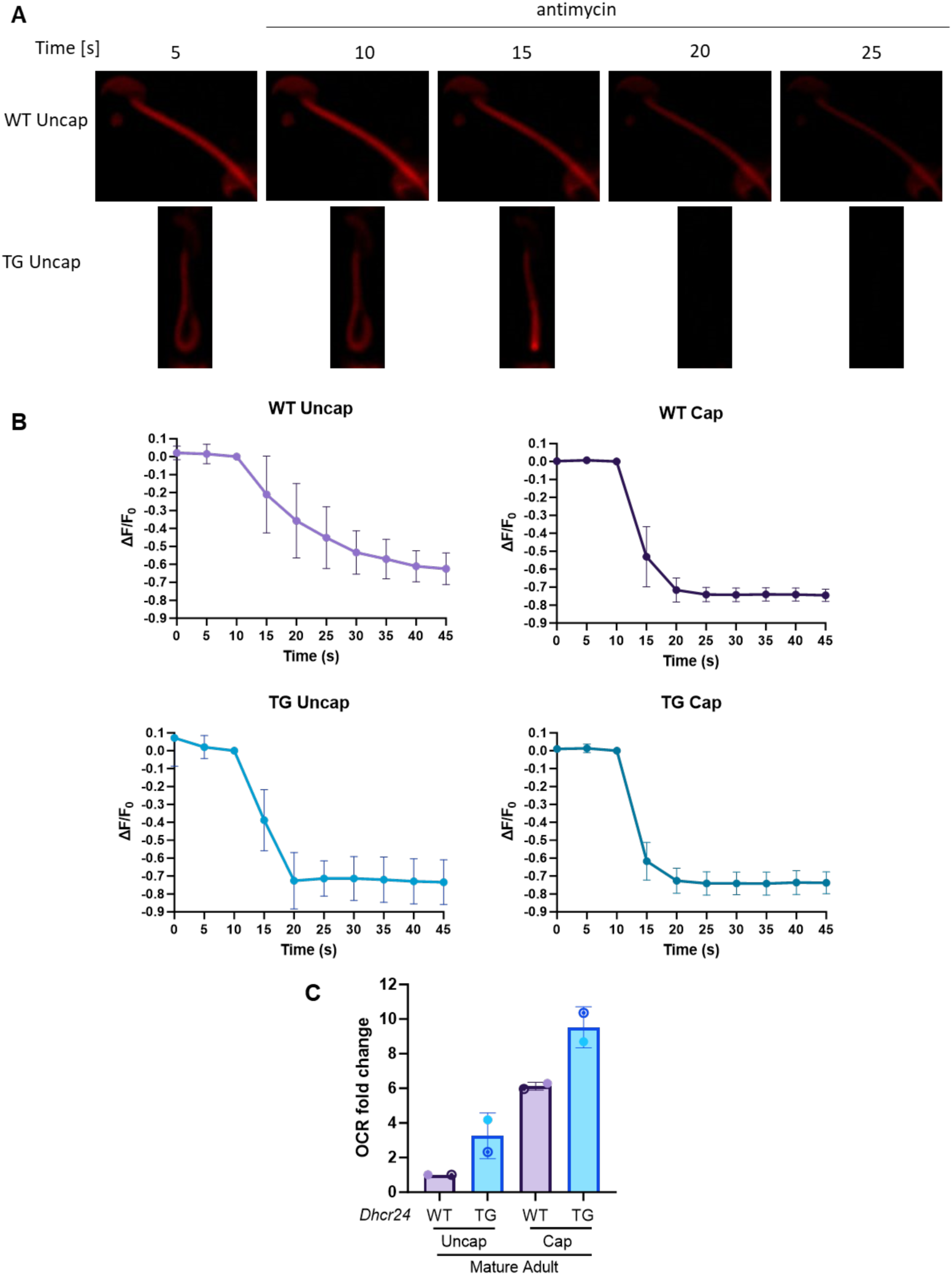
DHCR24 overexpression affects spermatozoan metabolism. (**A**) Dynamic representative images of mitochondrial membrane potential indicated by MitoTracker™ Red CMXRos in uncapacitated WT and *Dhcr24-TG* sperm upon antimycin treatment. (**B**) Transition of the mitochondrial membrane potential. The changes of fluorescence intensity were calculated as ΔF/F_0_ (F_0_, the mean fluorescence intensity of the sperm midpiece before adding antimycin A (t = 10 s); F, the fluorescence of the midpiece after adding antimycin A; ΔF=F− F_0_). Error bars SD, n=4. (**C**) Fold change in basal respiratory oxygen consumption rate from Seahorse analysis, measurements normalized to WT uncapacitated sperm. Spermatozoa from middle aged adult. Error bars SD.

In summary, desmosterol depletion and the consequent sterol imbalance results in an age-dependent decline in male fertility, partly due to reduced sperm count and motility with mitochondrial structural and functional defects.

## Discussion

### Dysregulation of lipid metabolism affects male fertility

Wechsler *et al*., (2003) previously reported that loss of DHCR24 function results in infertility in both female and male mice, with testicular abnormalities observed in *Dhcr24^-/-^*male mice. In this study, we report that gain of DHCR24 function, leading to desmosterol depletion, renders males subfertile and this condition worsens with age, while females remain fertile. Interestingly, an age-associated decline in male fertility has been reported in several other mouse models with dysregulated cholesterol homeostasis. Examples include ATP-binding cassette transporter 1 (ABCA1), a lipid transporter responsible for the efflux of excess cholesterol and phospholipids from Sertoli cells in the testis (Selva *et al*., 2004; Morales *et al*., 2008), and liver X receptor/retinoid X receptor (LXR/RXR) (Volle *et al*., 2007), which regulates ABCA1 expression (Venkateswaran *et al*., 2000; Costet *et al*., 2000). Consistently, lipid droplets accumulated in Sertoli cells of *Abca1*^-/-^ males, reducing their fertility by 6 months of age with decreased sperm count and testosterone levels (Selva *et al*., 2004). Similarly, accumulation of cholesterol/lipid droplets was observed in the testes of LXR-β-null males with age-dependent spermatogenesis defects (Robertson *et al.,* 2005). Since our previous study showed that desmosterol depletion in macrophages resulted in impaired LXR activation and lipid accumulation as well as reduced ABCA1 levels (Zhang et al., 2021^a^), we initially speculated that impaired LXR/RXR activation and/or ABCA1 expression might contribute to the phenotypes we observed in *Dhcr24-TG* males. However, our data comparing *Nr1h2* (LXRβ), *Nrhh3* (LXRα), and *Abca1* mRNA levels, as well as ABCA1 protein levels in the testes of WT versus TG animals showed no significant changes in either young or mature adult males (Figure 2–figure Supplement 2A and 2B). It is possible that desmosterol does not transcriptionally activates LXR in the mouse male reproductive system, or that desmosterol-regulated LXR signaling only manifests in animals fed a high fat diet and/or overweight animals, but these factors may not be responsible for the observed phenotype in our *Dhcr24-TG* mouse model. Similarly, desmosterol does not activate LXR in hepatocytes (Muse *et al*., 2018), suggesting that this intermediate of cholesterol biosynthesis is a specific LXR ligand in myeloid cells, but not in other cells and tissues (Muse *et al*., 2018; Spann *et al*., 2012; Zhang *et al*., 2021^a^).

Notably, DHCR24 overexpression in our mouse model had no observable effect on body fat mass (Figure 2–figure Supplement 3A) or plasma cholesterol levels (Figure 2–figure Supplement 3B). Consistently, there were only marginal changes in the immediate cholesterol precursors 7-DHC and 8,9-DHC in spermatozoa (Figure 2B), and no differences in the cholesterol levels isolated from the testis (Figure 2C). These findings are likely attributed to the negative feedback mechanisms that occur during spermatogenesis and spermiogenesis, ensuring the maintenance of appropriate cholesterol levels, given its crucial role. Furthermore, the normal fat to lean mass ratio in *Dhcr24-TG* mice rules out any known effects of obesity on male fertility, such as increased inflammation or abnormalities in hormone synthesis (Katib, 2015, Han et al., 2023). Instead, the observed defects result directly from an imbalance in cellular cholesterol homeostasis, specifically desmosterol depletion. A previous study indicated alterations in the desmosterol to cholesterol ratio in sperm from infertile men (Zalata et al., 2010). Thus, the sterol imbalance in the *Dhcr24-TG* spermatozoa itself is likely the cause of the delayed acrosome reaction. Additionally, the global defects in lipid signaling, levels of desmosterol, and other cholesterol precursors in the testis and plasma may affect spermatogenesis or spermiogenesis, contributing to the observed sperm defects and reduced sperm counts in *Dhcr24-TG* animals.

### Desmosterol-depleted sperm show defective cellular metabolism

We found abnormalities in the tail structure of sperm from *Dhcr24-TG* animals, particularly in the midpiece, as evidenced by mitochondrial packing defects and abnormally bent tail (Figure 4). It remains unclear whether these defects result from issues during spermatogenesis/spermiogenesis or are caused by changes in the lipid composition of the sperm plasma membrane and/or the mitochondrial membrane. Notably, these mitochondrial structural changes are associated with alterations in cellular metabolism in uncapacitated *Dhcr24-TG* spermatozoa. In previous studies, elevated OXPHOS levels are found in diabetic cardiac microvascular endothelial cells (CMECs), correlated with increased intracellular ROS (Zhang et al., 2021^b^). We previously showed that desmosterol depletion in macrophages also leads to ROS accumulation (Zhang et al., 2021^a^). In spermatozoa, mitochondria are a source of ROS with a dual function: ROS is a signaling molecule during capacitation (O’Flaherty et al., 2006, Boerke et al., 2013), but high levels of ROS have the potential to damage sperm DNA and negatively affect fertility (Aitken and Baker, 2006). Complex I and complex III are recognized as the major ROS producers in spermatozoa (Koppers et al., 2008). Our OXPHOS measurements revealed an increase in basal OCR levels in uncapacitated *Dhcr24-TG* spermatozoa, approaching levels similar to capacitated WT spermatozoa (Figure 5C). Furthermore, uncapacitated *Dhcr24-TG* sperm membranes are depolarized after antimycin treatment at rates similar to those observed in capacitated WT and *Dhcr24-TG* spermatozoa (Figure 5B). The sensitivity to antimycin in uncapacitated *Dhcr24-TG* spermatozoa and increased MMP could be explained by increased complex III activity, potentially leading to increased ROS production, which in turn may contribute to the observed sperm defects in TG spermatozoa.

In this study, we utilized a mouse model with DHCR24 overexpression to induce desmosterol deficiency, aiming to investigate significance of sterol homeostasis in sperm function and fertility, and explore the effects of sterols on sperm cell metabolism. Our findings revealed alterations in the sperm sterol composition, resulting in morphological and metabolic defects of spermatozoa, ultimately leading to impaired fertility. This underscores the impact of sterol imbalance on the fertilization ability of spermatozoa. Interestingly, we observed increased OCR and MMP. Whether *Dhcr24-TG* sperm accumulate more ROS as a result of sterol dysfunction remains to be investigated in the future. Importantly, our results establish a connection between cellular metabolism and sterol homeostasis, specifically highlighting the role of desmosterol. Furthermore, our studies suggest that deviations in sterol homeostasis may not always be correlated with obesity, at least in mice. If this finding extends to humans, it could have implications for how we understand and study sterol dysregulation and its relationship with infertility. Single nucleotide polymorphisms in genes encoding enzymes important for sterol synthesis may impact male fertility, even in the absence of increased body fat, while obesity in these individuals could worsen fertility outcomes.

## Materials and Methods

### Generation of *Dhcr24^fl/fl-CRE^* (*Dhcr24-TG*) knock-in mice

Homozygous *Dhcr24^fl/fl^* mice were crossed with the transgenic mice expressing Cre recombinase under the control of the EIIa promoter (JAX stock #003724) that targets expression of Cre recombinase to the early mouse embryo to induce germ line deletion of loxP flanked sites. *Dhcr24^fl/fl-CRE^* (*Dhcr24-TG*) mice were compared to *Dhcr24^fl/fl^*used as WT control group.

### Plasma cholesterol concentration measurements

Blood samples from young adult males were collected by retroorbital venous plexus puncture or cardiac puncture if terminal, after 6h fasting. Plasma was separated by centrifugation and stored at −80°C until analysis. Total plasma cholesterol levels were determined by standard enzymatic methods (Wako Chemicals).

### Antibodies and reagents

All antibodies and reagents used in this study are commercially available and listed in the key resource table. All the chemicals were from Sigma Aldrich unless indicated.

### Fertility test

#### Mating test

Wild type or *Dhcr24-TG* female mice were caged with biallelic *Dhcr24-TG* males for up to 4 months to record pregnancy and litter size when they gave births.

#### In vitro fertilization assay

The whole procedure was carried out using TYH media (in mM: 119.37 NaCl, 4.78 KCl, 1.19 KH_2_PO_4_, 5.56 glucose, 1 sodium pyruvate, 1.19 MgSO_4_, 25.07 NaHCO_3_, 1.71 CaCl_2_) supplemented with 0.0005% phenol red and 4 mg/mL of lipid rich BSA AlbuMax (Thermo Fisher). Young adult male mice were euthanized by CO_2_ and cervical dislocation. Sperm from one epididymis was released into TYH, and either capacitated for 1-1.5 hours without further dilution (final concentration 14-20×10^6^ sperm/mL) or diluted to concentration 2×10^6^ sperm/mL for capacitation. 7–10-week-old female B6D2F1 mice were superovulated by peritoneal injection of 10 IU PMSG 64 hours, and 10 IU hCG 16 hours prior to euthanasia by cervical dislocation. Cumulus-oocyte-complexes were isolated from ampullae of oviducts, placed into 50 µL TYH drop for 45 minutes - 1 hr. 10-20 µL of capacitated sperm were added to a TYH drops containing COCs, and left incubating for 6 hours, after which the fertilizing eggs were washed three times, discarding dying eggs, and incubated at 37°C 5% CO_2_ overnight. The next day, cleavage rate was scored based on the formation of 2-cell stage embryos.

### Single cell RNA-seq analysis

Existing single cell RNA-seq databases were analyzed as previously described (Hwang et al., 2021). The raw count matrices for human (GSE109037) and mouse (GSE109033) testis single cell RNA (scRNA)-seq datasets (Hermann et al., 2018) were downloaded from Gene Expression Omnibus (GEO) database. The downloaded raw count matrices were processed for quality control using the Seurat package (ver.3.2.3) (Stuart et al., 2019) with cells with <200 expressed features or >9000 (GSE109033) or 10 000 (GSE109037) expressed features, and >20% (GSE109037) or 25% (GSE109033) mitochondrial transcript fraction were excluded. The data was normalized by total expression, scaled and log transformed, and 2000 highly variable features was identified, followed by PCA to reduce the number of dimensions representing each cell. Statistically significant 15 PCs were selected based on the JackStraw and Elbow plots and provided as input for constructing a K-nearest-neighbors (KNN) graph based on the Euclidean distance in PCA space. Cells were clustered by the Louvain algorithm with a resolution parameter 0.1. Uniform Manifold Approximation and Projection (UMAP) was used to visualize and explore cluster data. Marker genes that define each cluster were identified by comparing each cluster to all other clusters using the MAST (Finak et al., 2015) provided in Seurat package. In order to correct batch effects among samples and experiments, the Harmony package was applied (ver.1.0) (Korsunsky et al., 2019) to the datasets. The Markov Affinitybased Graph Imputation of Cells (MAGIC) algorithm (ver.2.0.3) (van Dijk et al., 2018) was used to denoise and the count matrix and impute the missing data. In testis datasets from adult human, we identified 23 896 high quality single cells that were clustered into seven major cell types, namely spermatogonia, spermatocytes, early spermatids, late spermatids, peritubular myoid cell, endothelial cell, and macrophage. Similarly, we identified 30 268 high quality single cells from eight adult and three postnatal day 6 mouse testis tissue samples and subsequently defined 11 major cell populations.

### qRT-PCR

RNA was extracted from the testis using RNeasy Mini Kit (Qiagen) according to manufacturer’s instructions. Briefly, 20-30 mg of testis in 600 µL of RLT buffer was homogenized using 20G needle attached to a syringe. Samples were transferred into a microtube, centrifuged for 3 minutes 21000 g and supernatant was mixed with 600 µL of 70% ethanol. Mixture was transferred into column, spun down for 30 s at 8000 g and flow through was discarded. Column was washed with 700 µL RW1 buffer twice, followed by two washes with 500 µL RPE buffer, spun down empty to remove residual buffer, transferred into a clean Eppendorf tube and RNA was eluted in 30 µL of RNAse free water after incubation at 50°C for 1 minute. Next, 1 µg of RNA was treated with DNAse I (Amplification grade) (ThermoFisher), and cDNA was synthesized using iScript™ cDNA Synthesis Kit (Bio-Rad) according to manufacturer’s instructions. qRT-PCR was carried out using Power SYBR™ Green PCR Master Mix (Applied Biosystems) and samples were run on 96-well plates on a Bio-Rad CFX96 machine at 95°C for 3 minutes, and 40 cycles at 95°C for 5 seconds and 60°C for 15 seconds, followed by a dissociation step. Primer sequences were as follows: *Nr1h2*-F: 5’-CGTCCACCATTGAGATCATGT-3’; *Nr1h2*-R: 5’-GCGAGAACTCGAAGATGGGA-3’; *Nr1h3*-F: 5’-GCGCTTTGCCCACTTTACTG-3’; *Nr1h3*-R: 5’-CTCCAGAAGCATGACCTCGAT-3’; *Abca1-*F: 5’-GGAAGTTGGCAAGGTTGGTG-3’; *Abca1*-R: 5’-CGCCACTGTAGTTACTGGCA-3’; *β-actin*-F: 5’-CGCAGCCACTGTCGAGTC-3’; *β-actin*-R: 5’-GTCATCCATGGCGAACTGGT-3’. Relative quantification was carried out using ΔΔCt method, normalizing to the most efficient reaction, and relative to *β-actin* expression.

### Body composition analysis

Middle-aged adult (8-10 months) male mice were placed in EchoMRI-500 body composition analysis device (EchoMRI) (Dai Pra et al., 2022). Three replicate measurements of fat mass and lean mass for each animal were taken.

### Plasma cholesterol concentration measurements

Blood samples from young adult males were collected by retroorbital venous plexus puncture or cardiac puncture if terminal, after 6 hours of fasting. Plasma was separated by centrifugation and stored at −80°C until analysis. Total plasma cholesterol levels were determined by standard enzymatic methods (Wako Chemicals).

### Mouse sperm preparation and *in vitro* capacitation

Epididymal spermatozoa from adult male mice (young adult (YA): 2-4 months; mature adult (MA): 5-7 months; middle-aged adult (MDA): 8-10 months) (Flurkey et al., 2007) were collected by swim-out from cauda epididymis in HS medium (in mM: 135 NaCl, 5 KCl, 1 MgSO_4_, 2 CaCl_2_, 20 HEPES (pH 7.5), 5 glucose, 10 sodium lactate, 1 sodium pyruvate, pH 7.4 adjusted with NaOH, 320 mOsm/L), or EmbryoMax® M2 medium (EMD Millipore). To induce capacitation, 2×10^6^ sperm/mL were incubated in human tubular fluid (HTF) medium (EMD Millipore) at 37°C, 5% CO_2_ for 90 min. Sperm either under uncapacitating conditions (M2 or in HS) or under capacitating conditions (HTF) were treated with methyl-β-cyclodextrin (MBCD, 1.3 mg/mL) and polyvinyl alcohol (PVA, 1 mg/mL) for 1 hour at 37°C to examine the efficiency of cholesterol efflux and its effect on capacitation (acrosome reaction) rate. In experiments where protein levels from lysates from cauda, corpus and caput sperm were compared, caput sperm were purified using Percoll gradients as previously described (Krapf et al., 2012; Galan et al., 2021). Briefly, sperm suspension is layered over a Percoll gradient containing 45% Percoll/PBS in upper phase, and 90% Percoll/PBS in lower phase, and centrifuged at 650 g for 25 minutes. Interphase contained purified caput sperm, which were washed in 1x PBS and prepared for protein extraction as described below.

### Sperm motility analysis

Aliquots of sperm were placed in slide chamber (CellVision, 20 µm depth) and motility was examined on a 37°C stage of a Nikon E200 microscope under 10X phase contrast objective (CFI Plan Achro 10X/0.25 Ph1 BM, Nikon). Images were analyzed for spermatozoa by head tracing via computer-assisted sperm analysis (CASA, Sperm Class Analyzer version 6.3, Microptic, Barcelona, Spain). Images were recorded (40 frames at 50 fps) using CMOS video camera (Basler acA1300-200um, Basler AG, Ahrensburg, Germany) and analyzed by computer-assisted sperm analysis (CASA, Sperm Class Analyzer version 6.3, Microptic, Barcelona, Spain). Sperm motility (%) was quantified, and motion parameters including curvilinear velocity (VCL, in mm/s) and amplitude of lateral head displacement (ALH, in mM) were measured. Over 200 motile sperm were analyzed for each trial, with 3-4 biological replicates performed for each genotype, as denoted in figure legends.

### Flagellar waveform analysis

To tether sperm head for planar beating, non-capacitated or capacitated spermatozoa (2-3×10^5^ cells) from adult male mice were transferred to the fibronectin-coated 37°C chamber for Delta T culture dish controller (Bioptechs) filled with HS and HEPES-buffered HTF medium (H-HTF) (Chung et al., 2017). Flagellar movements of the tethered sperm were recorded for 2 s with 200 fps using pco.edge sCMOS camera equipped in Axio observer Z1 microscope (Carl Zeiss). All movies were taken at 37°C within 10 minutes after transferring sperm to the imaging dish. FIJI software (Schindelin et al., 2012) was used to generate overlaid images to trace waveform of sperm flagella as previously described (Wang et al., 2022).

### Sperm immunocytochemistry

Sperm were washed in PBS twice, attached on glass coverslips, and fixed with 4% paraformaldehyde (PFA) in PBS at room temperature (RT) for 10-15 minutes. Fixed samples were permeabilized using 0.1% Triton X-100 in PBS at RT for 10 minutes and blocked with 10% normal goat serum in PBS at RT for 1 hr. Cells were stained with DHCR24 (1:500, Cell Signaling Technology) or IZUMO (1:250, Bio-Academia) antibody in 10% goat serum in PBS at 4°C overnight. After washing in PBS, the samples were incubated with secondary antibody (1:1,000) in 10% goat serum in PBS at RT for 1 hr. Hoechst was used to counterstain nucleus. Immunostained samples were mounted with Prolong gold (Invitrogen) and cured for 24 h, or with VectaShield (Vector laboratories), followed by confocal imaging with Zeiss LSM710 using Plan-Apochrombat 63X/1.40 or alpha Plan-APO 100X/1.46 oil objective lens (Carl Zeiss).

### Protein extraction and western blotting

#### Preparation of testis lysate

Mouse testis lysate was prepared for western blotting as previously described (Hwang et al., 2019; Ded et al., 2020). In short, mouse testes were homogenized in 0.32M sucrose and centrifuged at 1000 g for 10 min at 4°C to remove cell debris and nuclei. 1% Triton X-100 in PBS containing protease inhibitor cocktail (cOmplete Mini, EDTA-free, Roche) was added to the cleared homogenates to make total testis lysate. The lysates were centrifuged at 4°C, 14,000 g for 30 min and the supernatant was used for the downstream experiments. For experiments measuring ABCA1 expression, testes were homogenized in 1x RIPA buffer containing protease inhibitor cocktail (cOmplete Mini, EDTA-free, Roche), centrifuged at 4°C, 14,000 g for 30 min and the supernatant was used for the downstream experiments.

#### Preparation of whole sperm lysate and solubilized protein extracts

Whole sperm protein was extracted as previously described (Chung et al., 2011; 2014; 2017). In short, mouse epididymal spermatozoa washed in PBS were directly lysed in 2× SDS or 2x LDS sample buffer. The whole sperm lysate was centrifuged at 15,000 g, 4°C for 10 min. After adjusting DTT to 50 mM, supernatant was denatured at 75°C for 10 min before loading to gel. Antibodies used for western blotting were rabbit anti-mouse DHCR24 (1:1000, Cell Signaling Technologies), and anti-acetylated tubulin (1:20,000, Sigma). Secondary antibodies were anti-rabbit IgG-HRP (1:10,000) and anti-mouse IgG-HRP (1:10,000) from Jackson ImmunoResearch (West Grove, PA).

### Sperm lipidomics

#### Preparation of sperm samples

Sterol biosynthesis in sperm was analyzed by liquid chromatography with tandem mass spectrometry (LC-MS/MS) as described previously (Mitsche et al., 2015; Zhang et al., 2021^a^). Sperm cells from mature adult (5-7 months old) WT or *Dhcr24-TG* mice were isolated from cauda epididymis. Cells were washed with PBS twice, then snap frozen in liquid N2 and stored at −80 °C.

#### LC-MS/MS analysis

Lipids were extracted from the samples by a modified Bligh–Dyer extraction, and sterols were analyzed by LC-MS/MS as described previously (Zhang et al., 2021^a^). Briefly, lipid extracts from samples were dried under nitrogen and reconstituted in 90% methanol. Sterols were analyzed using a Shimadzu LC20ADxr high-performance liquid chromatograph equipped with an Agilent Poroshell 120 EC-C18 column (2.1 × 150 mm, 2.7-μm particles; Agilent Technologies). The elution was done using a solvent gradient that transitioned linearly from 93% methanol/7% H2O to 100% methanol in 7 min. The column was washed for 5 min in 100% methanol and then returned to the initial solvent. Sterols were detected using a SCIEX API 5000 triple quadrupole mass spectrometer equipped with a Turbo V APCI source in positive mode with atmospheric pressure chemical ionization at a temperature of 350 °C. Data were acquired under multiple reaction monitoring for mass pairs optimized for each sterol. The internal standards were commercially available for all but four of the sterols (dihydro-ff-MAS, dihydro-t-MAS, dehydrolathosterol, and dehydrodesmosterol) in the cholesterol biosynthetic pathway that were identified by their unique m/z values and retention times as described (Mitsche et al., 2015; McDonald et al., 2012).

### Scanning electron microscopy

Sperm cells were attached on the glass coverslips and fixed with 2.5% glutaraldehyde (GA) in 0.1 M sodium cacodylate buffer (pH 7.4) for one hour at 4°C and post fixed in 2% osmium tetroxide in 0.1 M cacodylate buffer (pH 7.4). The fixed samples were washed with 0.1 M cacodylate buffer for 3 times and dehydrated through a series of ethanol to 100%. The samples were dried using a 300 critical point dryer with liquid carbon dioxide as transitional fluid. The coverslips with dried samples were glued to aluminum stubs and sputter coated with 5 nm platinum using a Cressington 208HR (Ted Pella) rotary sputter coater. Prepared samples were imaged with Hitachi SU-70 scanning electron microscope (Hitachi High-Technologies).

### Transmitted electron microscopy

Collected epididymal sperm cells were washed and pelleted by centrifugation and fixed in 2.5% glutaraldehyde and 2% PFA in 0.1 M cacodylate buffer pH 7.4 for one hour at RT. Fixed sperm pellets were rinsed with 0.1 M cacodylate buffer and spun down in 2% agar. The chilled blocks were trimmed, rinsed in the 0.1 M cacodylate buffer, and replaced with 0.1% tannic acid in the buffer for one hour. After rinsing in the buffer, the samples were post-fixed in 1% osmium tetroxide and 1.5% potassium ferrocyanide in 0.1 M cacodylate buffer for one hour. The post-fixed samples were rinsed in the cacodylate buffer and distilled water, followed by en bloc staining in 2% aqueous uranyl acetate for one hour. Prepared samples were rinsed and dehydrated in an ethanol series to 100%. Dehydrated samples were infiltrated with epoxy resin Embed 812 (Electron Microscopy Sciences), placed in silicone molds, and baked for 24 hours at 60°C. The hardened blocks were sectioned in 60 nm thickness using Leica UltraCut UC7. The sections were collected on grids coated with formvar/carbon and contrast stained using 2% uranyl acetate and lead citrate. The grids were imaged using FEI TecnaiBiotwin Transmission Electron Microscope (FEI, Hillsbroro, OR) at 80 kV. Images were taken using MORADA CCD camera and iTEM (Olympus) software.

### Mitochondrial membrane potential (ΔΨm) measurement

1 million uncapacitated sperm from cauda epididymis in 500 µL HS were attached on Delta T chamber coated with poly-D-Lysine (0.1 mg/mL). 100 nM of Mitotracker Red CMXRos (ThermoFisher), previously established for MMP measurements (Pendergrass et al., 2004) including in sperm (Santiani et al., 2016), was loaded into sperm for 30 min, followed by a wash with HS. Alternatively, sperm were capacitated for 1 hour in HTF, then HTF was replaced with H-HTF, sperm were attached on poly-D-Lysine coated Delta T chamber and Mitotracker was loaded for 30 minutes, then washed with H-HTF. 10 µM of antimycin was added while taking time lapse images. The images were captured by Axio observer Z1 microscope (Carl Zeiss) equipped by pco.edge cMOS camera and DG-4. The data were analyzed by Zen Blue.

### Oxygen consumption rate measurement

For extracellular flux measurement, HTF was replaced by H-HTF for capacitated sperm and M2 by HS media for uncapacitated sperm. 2 million sperm per well was placed into concanavalin A (0.05 mg/mL) coated 96-well plate for Seahorse XFe96 (Agilent). Mitochondrial activity (OXPHOS) was measured after sequential injections of 5 µM oligomycin, 1 µM FCCP, and 1 µM of rotenone and antimycin A. Three measurements were taken after each injection. Basal respiration levels were calculated by subtracting oxygen consumption rate (OCR) values obtained after rotenone and antimycin injection from initial OCR measurements prior to injections (Balbach et al., 2020).

### Quantification and statistical analysis

Statistical analyses were performed using Student’s t-test or one-way analysis of variance (ANOVA) with Tukey or Bonferroni post hoc test using Graphpad Prism 9. Differences were considered significant at *p < 0.05, **p < 0.01, and ***p < 0.001.

### Key resource table

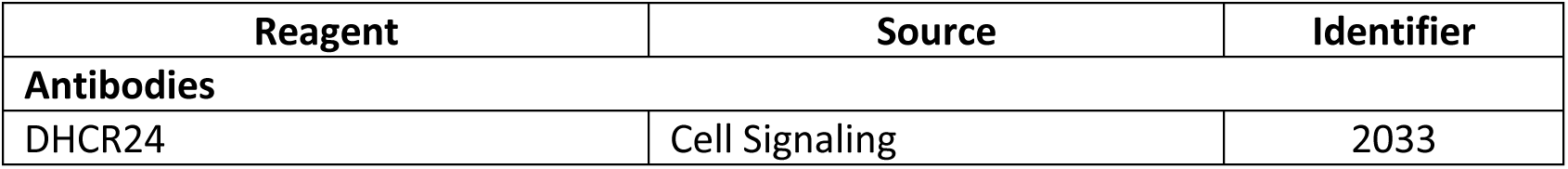
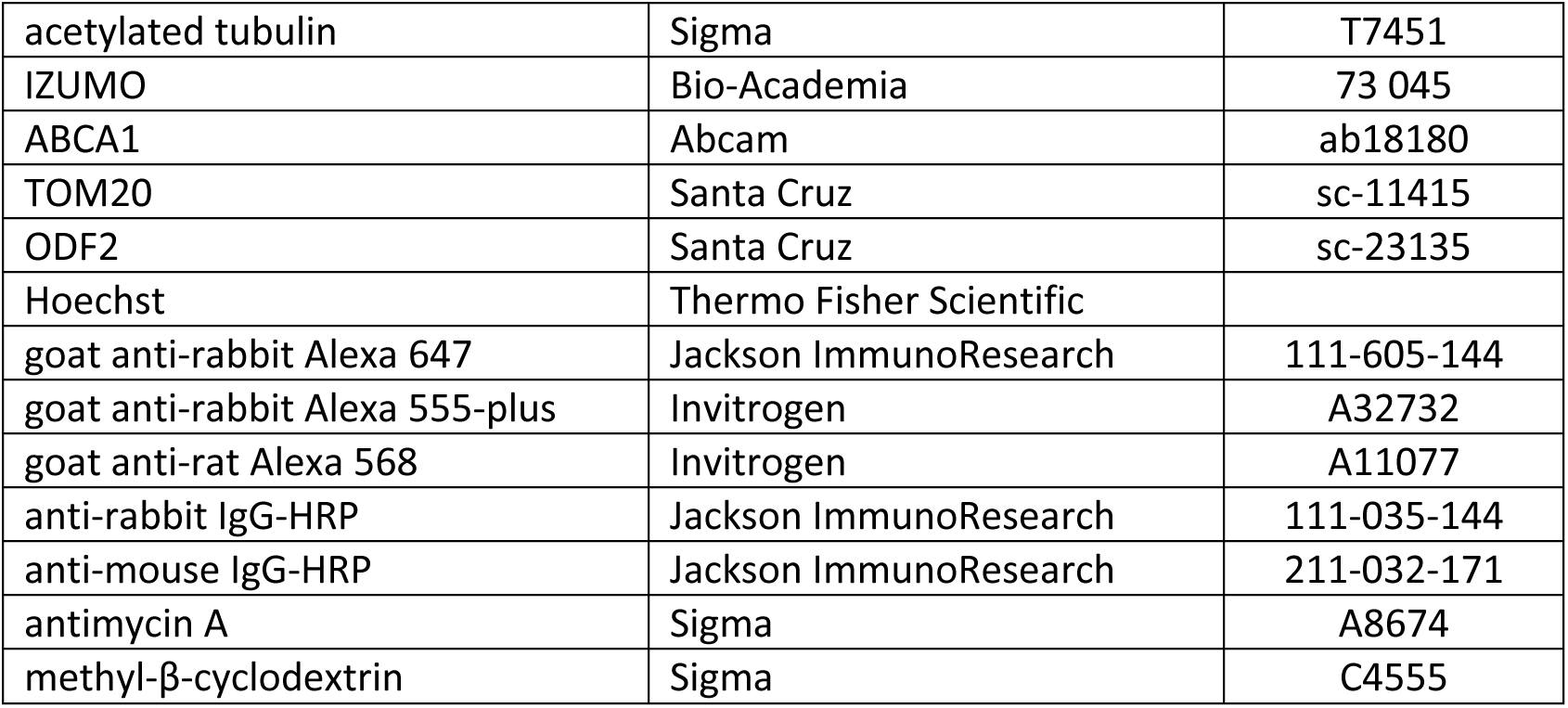

## Supporting information

Supplemental_Relovska et al

## Acknowledgements

We thank the Yale Center for Cellular and Molecular Imaging for assistance in electron microscopy for the SEM and TEM images, Madison Gowett for sperm immunocytochemistry, Olga Blanco Prieto for technical advice with Seahorse experiments, Rafael Dai Pra and Viktor Feketa for assisting and technical advice with EchoMRI, and Elena Gracheva for sharing EchoMRI.

## Additional Information

## Author contributions

Sona Relovska – Validation, Data Curation, Formal Analysis, Investigation, Visualization, Methodology, Writing – Original Draft Preparation, Writing – Review & Editing; Huafeng Wang - Validation, Data Curation, Formal Analysis, Investigation, Visualization, Methodology, Writing – Original Draft Preparation; Xinbo Zhang – Validation, Conceptualization, Data curation, Investigation; Kyung Jo Jeong – Formal Analysis, Software, Visualization; Jungmin Choi – Methodology, Visualization; Pablo Fernandez Tussy – Validation, Data Curation, Investigation, Methodology; Yajaira Suárez – Methodology, Resources, Writing – Review & Editing; Jeffrey G. McDonald – Validation, Data Curation, Formal Analysis, Methodology; Carlos Fernandez-Hernando - Conceptualization, Writing – Review & Editing, Supervision, Resources, Funding Acquisition; Jean-Ju Chung - Conceptualization, Data Curation, Formal Analysis, Investigation, Visualization, Methodology, Writing – Original Draft Preparation, Writing – Review & Editing, Project administration, Supervision, Resources, Funding Acquisition

## Ethics

Animal experimentation: This study was performed in strict accordance with the recommendations in the Guide for the Care and Use of Laboratory Animals of the National Institutes of Health. All the mice were treated in accordance with guidelines approved by Yale (11577) Animal Care and Use Committees (IACUC).

