## Supplemental_Relovska et al for "DHCR24-mediated sterol homeostasis during spermatogenesis is required for sperm mitochondrial sheath formation and impacts male fertility over time"

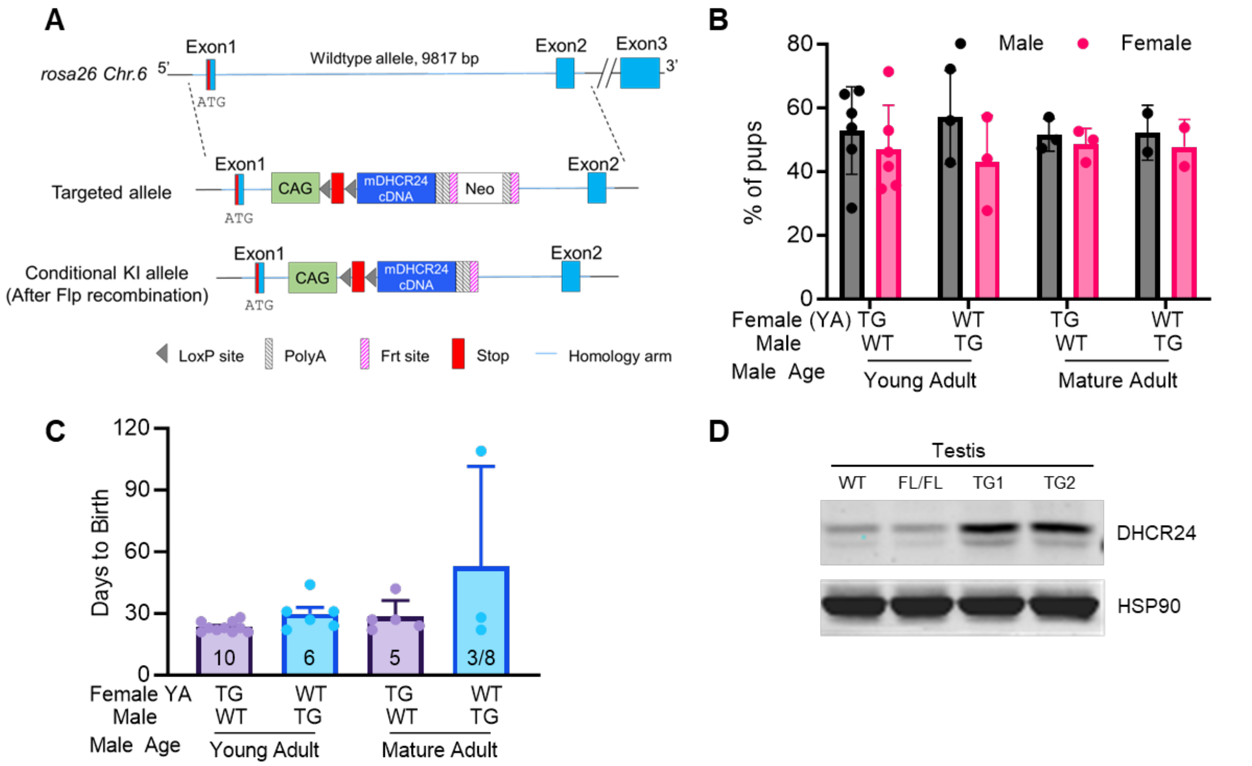


**Figure supplement 1.** (**A**) Genomic structures of targeted alleles for *Dhcr24-TG* mice generation. (**B**) Sex ratios of pups from wild-type and *Dhcr24-TG* young and mature adult mice mated with young adult females. (**C**) Number of days to birth from wild-type and *Dhcr24-TG* young and mature adult male mice mated with young adult females. (**D**) DHCR24 protein levels in testis from wild type, floxed (FL/FL), and *Dhcr24-TG* young adult animals. HSP90 used as a control. All error bars SD.

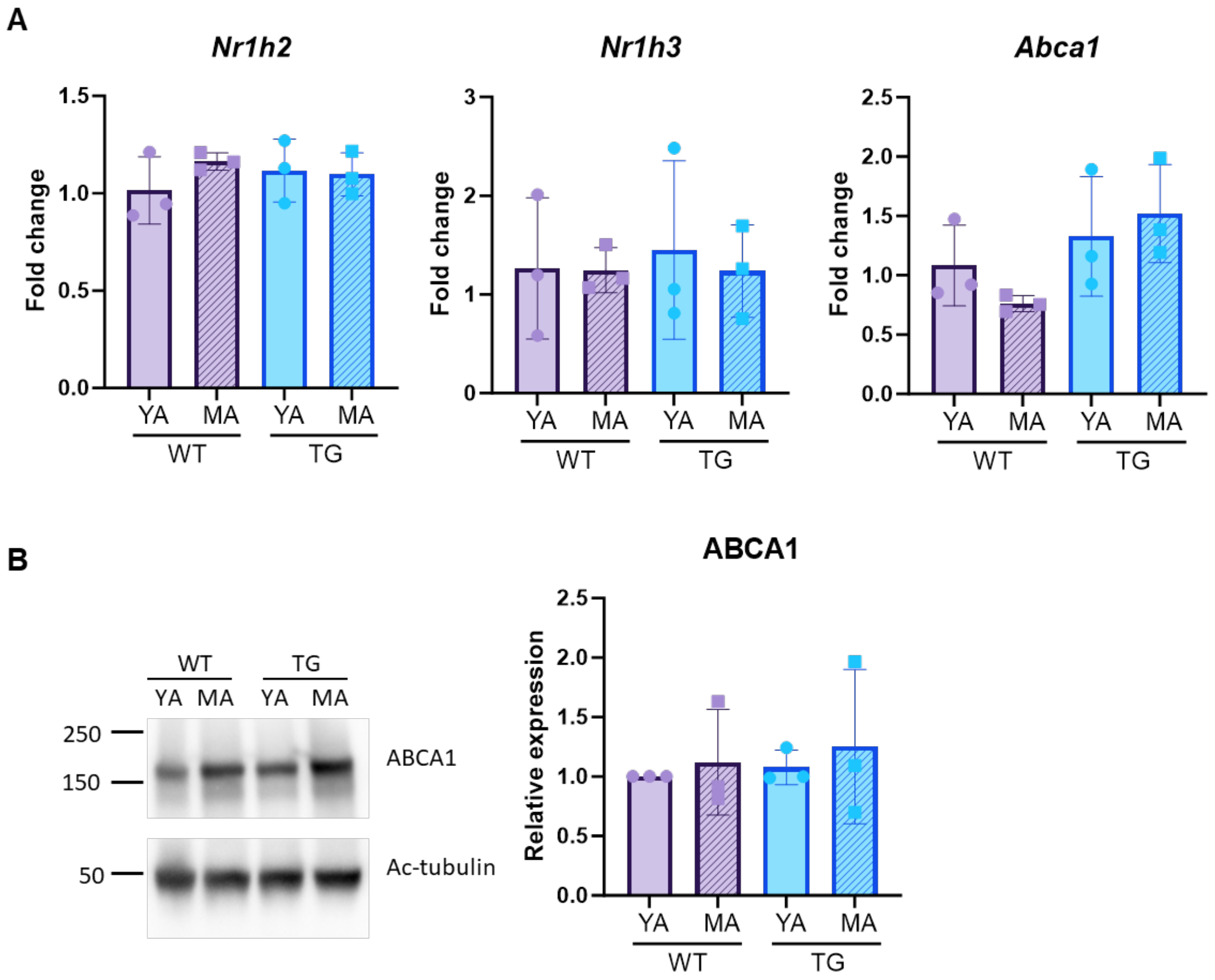


**Figure supplement 2.** (**A**) Quantification of *Nr1h2*, *Nr1h3* and *Abca1* mRNAs by qRT-PCR. *Nr1h2* and *Nr1h3* encode LXRβ and LXRα, respectively. *β-actin* is used as a housekeeping control gene. (**B**) Western blot (*left*) and quantification of ABCA1 protein levels normalized by acetylated tubulin in the testis from WT and *Dhcr24-TG* young and mature adult male mice (*right*). Statistical analysis one way ANOVA. YA, young adult (2 – 4 months old); MA, mature adult (5 – 7 months old). All error bars SD.

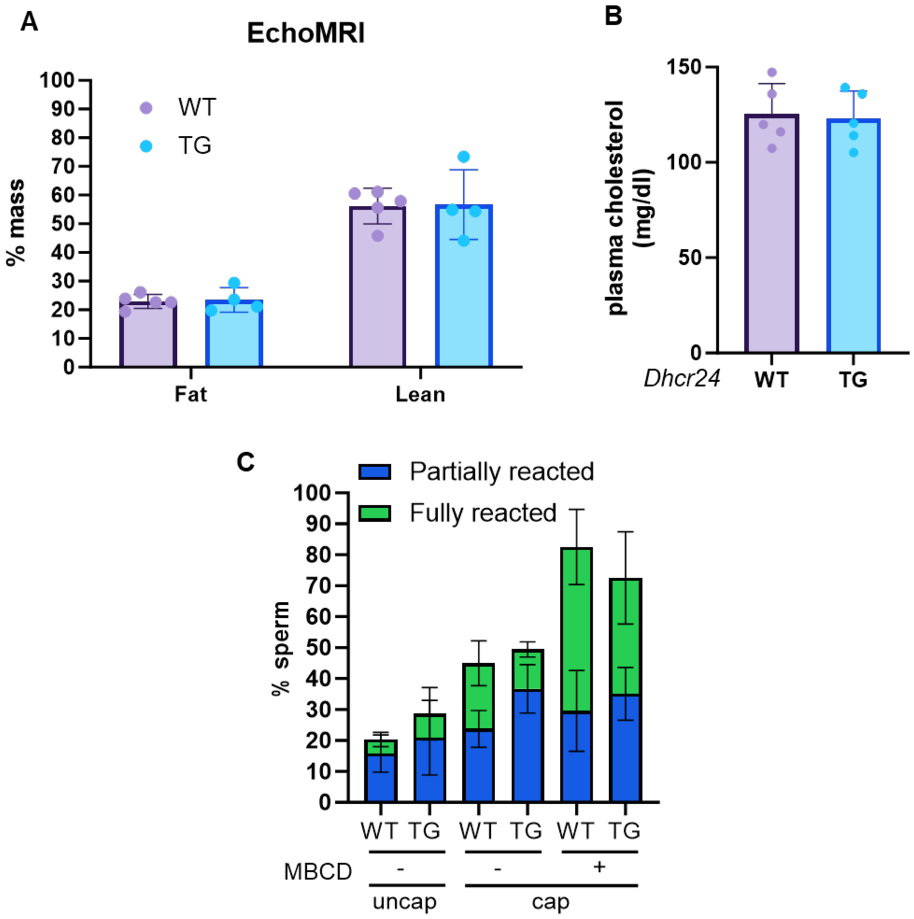


**Figure supplement 3.** (**A**) Percentage of lean and fat mass in middle aged adult WT and *Dhcr24-TG* (TG) animals measured by EchoMRI. (**B**) Plasma cholesterol levels of WT and *Dhcr24-TG* young adult animals (**C**) Acrosome reaction progression by IZUMO staining. Spermatozoa from middle aged adult WT and *Dhcr24-TG* (TG) animals were incubated under non-capacitating (uncap) or capacitating (cap) conditions in the absence (-) or presence (+) of methyl-β-cyclodextrin (MBCD). All error bars SD.

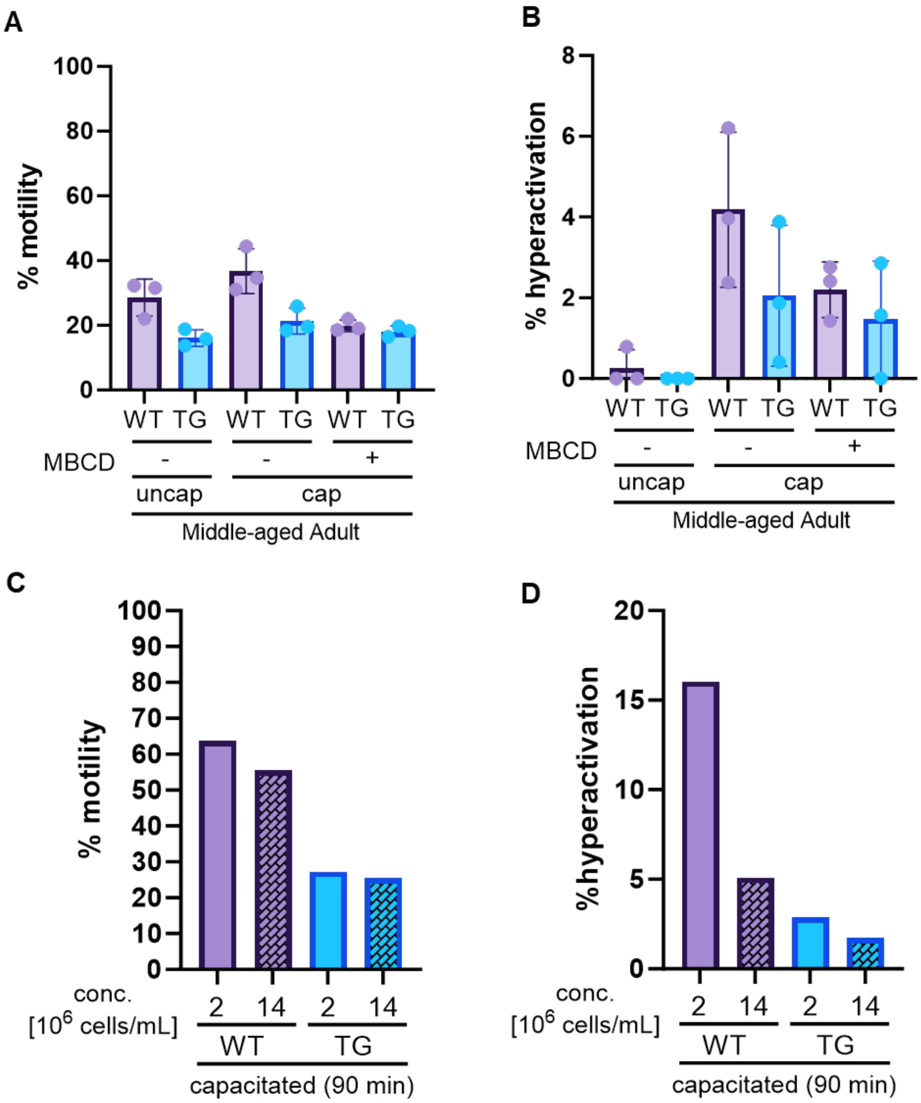


**Figure supplement 4.** (**A** and **B**) Motility (A) and hyperactivated motility (B) of spermatozoa from middle aged adult WT and *Dhcr24-TG* (TG) animals incubated under non-capacitating (uncap) or capacitating (cap) conditions in the absence (-) or presence (+) of methyl-β-cyclodextrin (MBCD). (**C** and **D**) Total motility (C) and hyperactivated motility (D) of sperm from WT and *Dhcr24-TG* (TG) animals capacitated at high ($14x10^6$ sperm/mL) and standard ($2x10^6$ sperm/mL) concentration. All error bars SD.

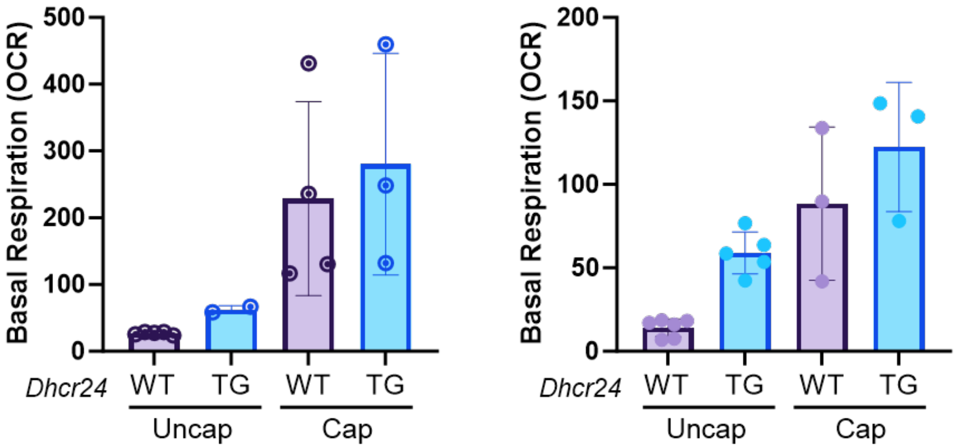


**Figure supplement 5.** Basal respiratory OCR of uncapacitated and capacitated sperm from middle aged adult WT and *Dhcr24-TG* (TG) animals. Data from Seahorse analysis calculated as initial measurements to which measurements after antimycin/rotenone treatment were subtracted. Graphs show two biological replicates, and each dot represents technical replicates. All error bars SD.
